# A synthetic MALDI-TOF MS atlas for microbial classification across hospitals

**DOI:** 10.64898/2026.10.19.759651

**Authors:** Alejandro Guerrero-López, Eline Meijer, Yukino Gütlin, Kristoffer Koch, Nighat Rabbani, Eva Hitz, Esther Serrano-Conde, Benjamin P. Howden, Kirstine K Søgaard, Muhammad Ali Syed, Michael Oberle, Jacob Moran-Gilad, Marta Hernández-García, Norelle L. Sherry, Adrian Egli

**Affiliations:** Institute for Medical Microbiology, University of Zurich, Zurich, Switzerland; Department of Clinical Microbiology, Aalborg University Hospital, Aalborg, Denmark; Department of Clinical Medicine, Aalborg University, Aalborg, Denmark; Department of Microbiology, The University of Haripur, Hattar Road, 22620, Haripur, Pakistan; Institute of Laboratory Medicine, Kantonsspital Aarau, 5001, Aarau, Switzerland; Servicio de Microbiología, Hospital Ramón y Cajal and Instituto Ramón y Cajal de Investigación Sanitaria (IRYCIS), Madrid, Spain; Microbiological Diagnostic Unit Public Health Laboratory, Department of Microbiology and Immunology, University of Melbourne at the Peter Doherty Institute for Infection & Immunity, Melbourne, Victoria, Australia; Centre for Pathogen Genomics, University of Melbourne, Melbourne, Victoria, Australia; Department of Infectious Diseases & Immunology, Austin Health, Melbourne, Victoria, Australia; Department of Microbiology, Royal Melbourne Hospital, Melbourne, Victoria, Australia; Clinical Microbiology Laboratory, Department of Clinical Microbiology and Infectious Diseases, Hadassah Medical Center, Jerusalem, Israel; CIBER de Enfermedades Infecciosas (CIBERINFEC), Instituto de Salud Carlos III (ISCIII), Madrid, Spain

## Abstract

Matrix-Assisted Laser Desorption/Ionization Time-of-Flight (MALDI–TOF) mass spectrometry (MS) is central to routine clinical microbiology for species identification. Its combination with machine learning is constrained by fragmented public datasets, species imbalance and acquisition-related variability. Synthetic datasets could provide more training data, but they must preserve the spectral information needed to make predictions on real samples.

Here, we present an open, synthetic MALDI–TOF MS atlas comprising 4.3 million spectra across 43 clinically relevant bacterial species. To construct the atlas, we harmonised public repositories from three countries and trained a species- and institution-conditioned concept-bottleneck generative variational autoencoder (CBGM-VAE) on 179,086 measured spectra.

We assessed the atlas as both a stand-alone training resource and a complement to real data, using bacterial species classification as a functional benchmark. Evaluation comprised 7,269 real spectra from more than 7 countries, 24 institutions and multiple acquisition platforms, covering: temporal, technical, biological and combined shifts. A neural network trained only on synthetic spectra achieved F1 scores ranging from 0.788 to 1.000 across external cohorts. When synthetic spectra were combined with real training data, classification performance improved or matched using real data only.

These findings demonstrate that the atlas retains transferable information and can complement measured data. The openly released atlas provides a reusable resource for supervised learning and benchmarking. It can also be explored for pretraining representation-learning and foundation models in MALDI–TOF MS research.

## Introduction

Matrix-Assisted Laser Desorption/Ionization Time-of-Flight mass spectrometry (MALDI–TOF MS) is the reference standard for pathogen identification in clinical microbiology routine diagnostics. MALDI-TOF MS allows rapid identification at the species level for more than 4,500 species from single colonies within minutes [1]. Moreover, there is a high potential for MALDI-TOF MS beyond its broadly applied routine use in species identification. In combination with machine learning, spectral signatures could be associated with subspecies typing [2], detection of antimicrobial resistance [3], and virulence [4]. Yet, these applications remain limited by strong taxonomic imbalance, fragmented public datasets, and substantial inter-laboratory variability [5, 6]. Benchmarking studies have shown that differences in instruments, protocols, and acquisition settings can affect external generalization [7].

Generative modelling offers a promising strategy to expand the available MALDI– TOF MS training data. They could synthesise spectra and organise them into balanced, shareable resources for model development. Such resources could be used either as stand- alone training datasets or in combination with real spectra. However, the usefulness of synthetic spectra depends on whether they preserve the biological information needed for downstream tasks and provide additional value beyond already publicly available data. Lately, data-hungry self-supervised models are emerging in the MALDI-TOF MS research field [6, 8–10]. However, current augmentation techniques treat spectra as only species-conditioned signals and do not explicitly model acquisition context [11, 12], despite site-, instrument-, and protocol-specific variations being major drivers of domain shift and a central barrier to robust diagnostic deployment across laboratories [5–7].

Here, we built an open, species-balanced synthetic MALDI–TOF MS atlas comprising 4.3 million spectra across 43 clinically relevant bacterial species. To construct the atlas, we curated public MALDI–TOF MS repositories from three countries and six institutions. We defined a set of species based on the pathogen groups defined in the Lancet Global Burden of Disease Study [13]. After expanding the list to match the species available in the public repositories, we obtained 43 bacterial species and 263,618 measured spectra. Of these, 179,086 spectra were used to train a Concept Bottleneck Generative Model Variational Autoencoder (CBGM-VAE), conditioned on both species and institution (Fig. 1). Institution was included as a proxy for differences in acquisition conditions.

**Fig. 1:**
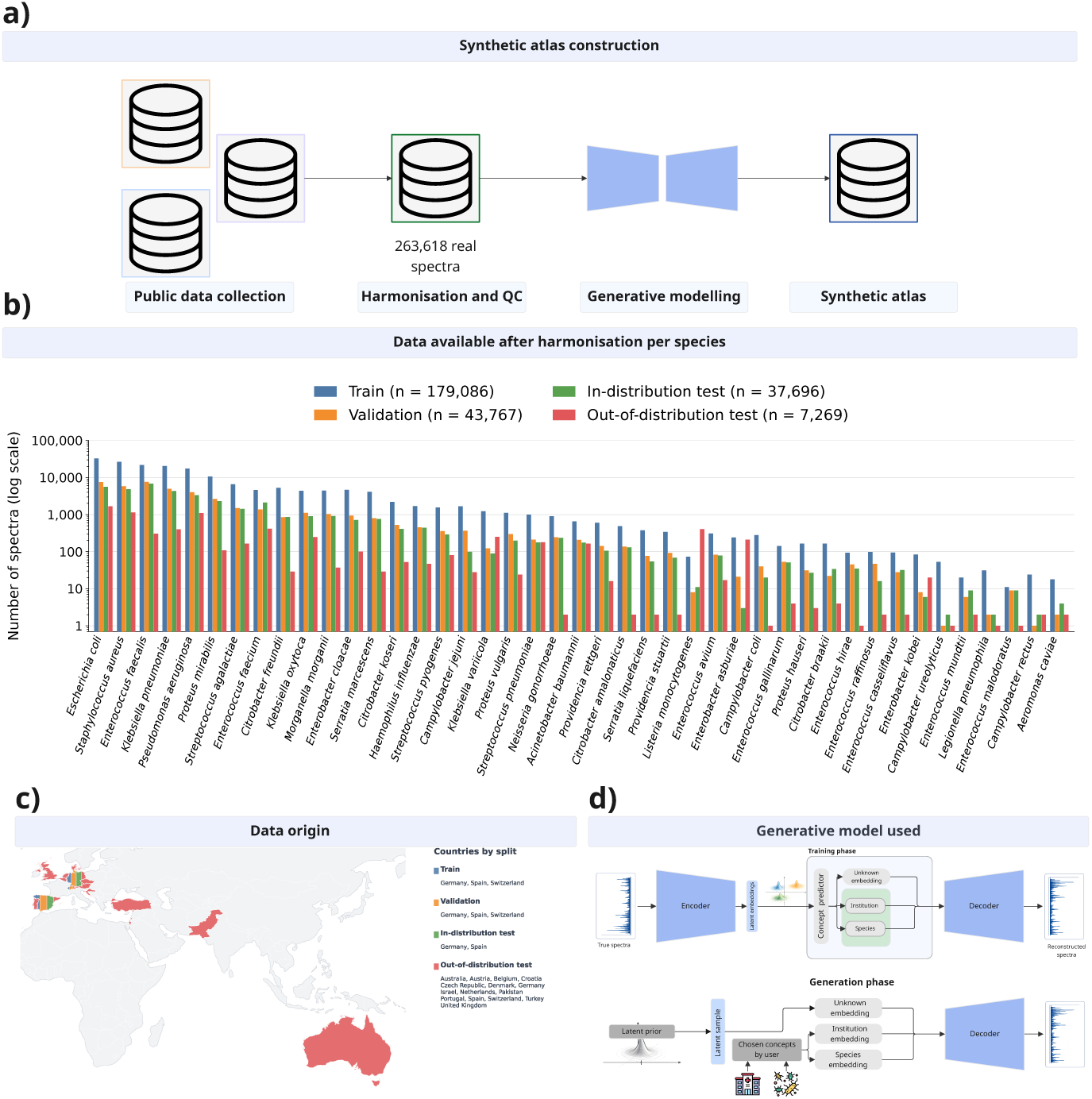
Synthetic atlas creation. **a,** Synthetic atlas diagram construction, from data collection to generative modelling. **b,** Species-level composition of all data used per species after harmonisation. **c,** Geographic origin of all spectra used, coloured by split role. **d,** Architecture of the CBGM-VAE used to construct the synthetic atlas. Specific details on quality control, data harmonisation, and dataset splitting (train, validation and test) are provided in the Methods section.

We evaluated the atlas as both a stand-alone training resource and a complement to real data, using bacterial species classification as a benchmark. The external panel contained 7,269 real spectra. Of these, 4,971 originated from 24 institutions across seven countries. The remaining 2,298 came from the multicentre study by Cúenod et al. [7], which covered 36 institutions across 12 countries. The evaluation included multiple acquisition platforms and tested performance under temporal, technical, biological, and combined shifts. A classifier trained only on the atlas identified species in real external samples. Combining synthetic and real training data gave improvements in weighted F1 over real-only training in several cohorts and matched the reported scores in the rest. By releasing the atlas openly [14], we provide a reusable resource for supervised learning and benchmarking, and for exploring representation learning and foundation-model pretraining in MALDI–TOF MS.

## Results

### An open synthetic MALDI–TOF MS atlas

We constructed an open synthetic MALDI–TOF MS atlas containing **4.3 million spectra**, with 100,000 synthetic spectra for each of the 43 species analysed in this study.

To construct the atlas, we first harmonised publicly available MALDI–TOF MS data, as shown in Fig. 1a. Specifically, we curated three public resources spanning six institutions across three countries: Database of Resistance Information on Antimicrobials and MALDI-TOF MS (DRIAMS) (four institutions, A to D, Switzerland) [5], MARISMa (one institution, E, Spain) [15], and MS-UMG (one institution, F, Germany) [16]. All institutions in this manuscript are renamed from A to AB, e.g., Institution A or Institution K. Their full names and corresponding identifiers are provided in the Supplementary Institution naming and standardisation section.

We based the species selection on the bacterial pathogen groups reported in the Lancet Global Burden of Disease Study [13]. Because several groups were defined at the genus level, we expanded these definitions to the species represented in the public databases, resulting in 43 bacterial species as shown in Fig. 1b. After quality control and harmonisation, the combined dataset contained 263,618 measured spectra (Fig. 1a), of which 179,086 were used for generative model training as shown in Fig. 1b.

Harmonised data contained an acquisition-related structure, as shown in Supplementary Fig. S1. Spectra clustered primarily by species but also showed structure associated with institution, database, and country, consistent with previous research using these databases [5, 6].

To capture both taxonomic and acquisition-related variation, we used a Concept Bottleneck Generative Model Variational AutoEncoder (CBGM-VAE) [17] as the generative model. A CBGM-VAE can incorporate multiple conditioning concepts associated with each input sample. For this purpose, we used species and institution to condition our spectra. Intuitively, species captures the dominant taxonomic structure of the spectrum, whereas institution acts as a proxy for acquisition-related effects arising from differences in epidemiology, instruments, protocols, sample preparation, and local workflows. This granular data is missing for public databases; therefore, the institution is the most granular information available.

The atlas is published openly for the research community for training and pretraining purposes [14].

### The synthetic atlas preserves spectral and species structure

We first tested whether the synthetic atlas showed the spectral and taxonomic structure observed in real MALDI–TOF MS data.

#### Qualitative comparison of real and synthetic spectra

First, we qualitatively compared real and synthetic spectra for five important species: *Acinetobacter baumannii*, *Enterococcus faecium*, *Escherichia coli*, *Pseudomonas aeruginosa* and *Staphylococcus aureus* in Institutions E and F (Fig. 2). The left panels show the average measured spectrum for each species at each institution, with Institution F displayed in the upper half and Institution E in the lower half. The right panels show the same measured spectra, now overlaid with the average synthetic spectra generated for the corresponding species and conditioned on each institution. This allows a direct comparison between measured and synthetic spectra while preserving the differences between institutions.

**Fig. 2:**
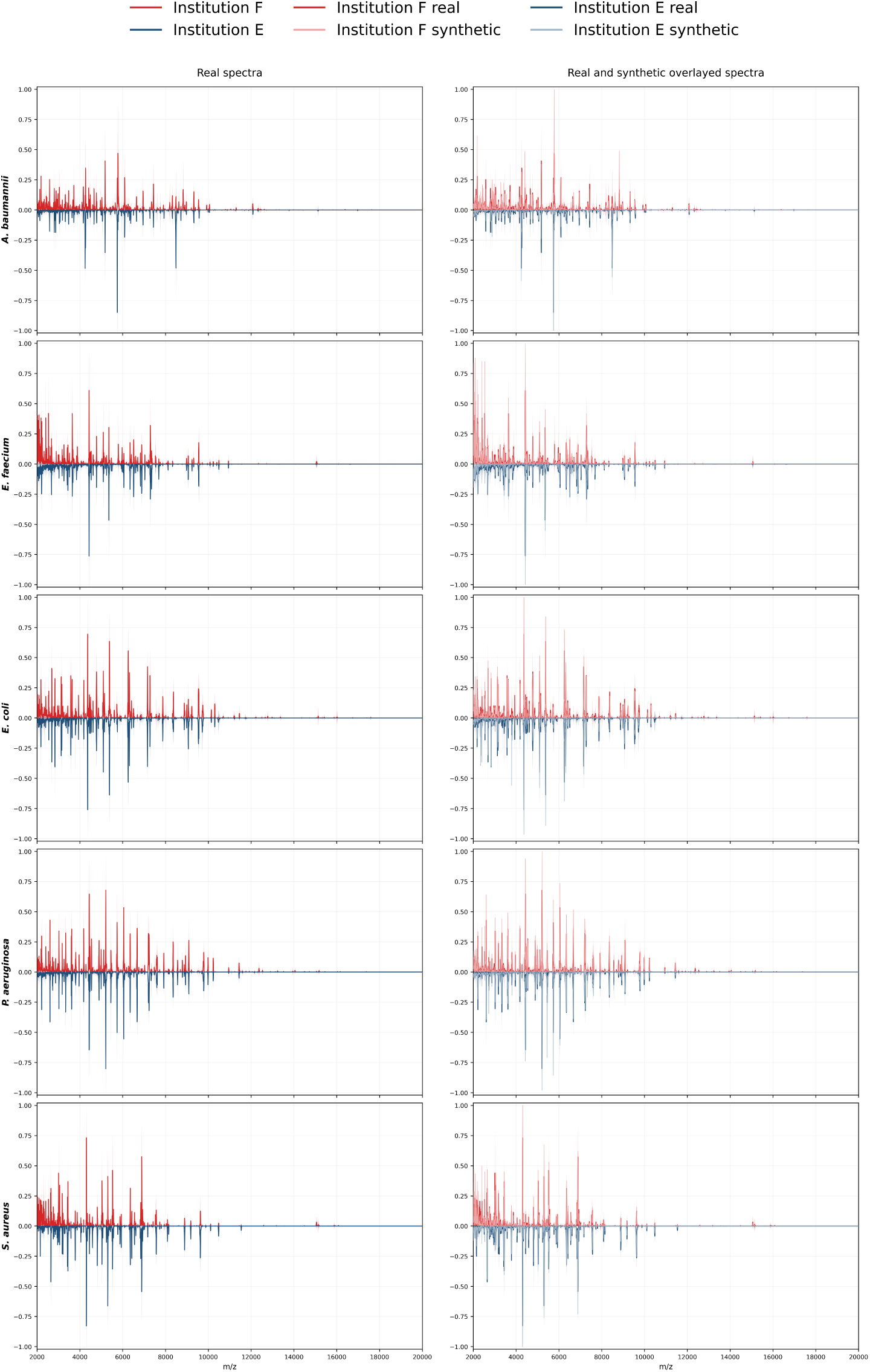
Synthetic spectra preserve species- and institution-associated MALDI–TOF MS structure. Rows show five representative species and columns separate measured spectra (left) from measured and synthetic overlays (right). Colours denote institution: red for Institution F and dark blue for Institution E. Solid lines show measured mean spectra, dashed lines show synthetic mean spectra, and shaded bands indicate ±1*σ* of both distributions. A zoom in can be found in Supplementary Fig. S2.

Across species, the synthetic spectra reproduced the dominant peak positions and overall intensity envelopes observed in the measured data. The generated spectra also retained visible institution differences, as seen in Fig. 2b. Synthetic spectra have different peak distribution per institution. This suggests that atlas generation preserved, qualitatively, both dominant taxonomic signal and part of the acquisition-related variability present in the source data.

This structure was also kept in the complete dataset level. We jointly projected measured and synthetic spectra using t-SNE (Fig. 3). Synthetic and measured spectra formed coherent species-specific clusters (Fig. 3a), indicating that the generated data retained the broad taxonomic organisation of the measured spectra. When coloured by institution, synthetic spectra appeared closer to real spectra from the same institution (Fig. 3b). This was observed when the model was conditioned to generate spectra from a specific institution. This is visible as in each Institution cluster, both measured (points) and synthetic samples (crosses) are found. In contrast, spectra generated using uniform-institution (equi-institution) sampling remained grouped by species (grey in Fig. 3b). However, they were less closely associated with specific institutions and appeared to cover a broader range of spectral variation within each species.

**Fig. 3:**
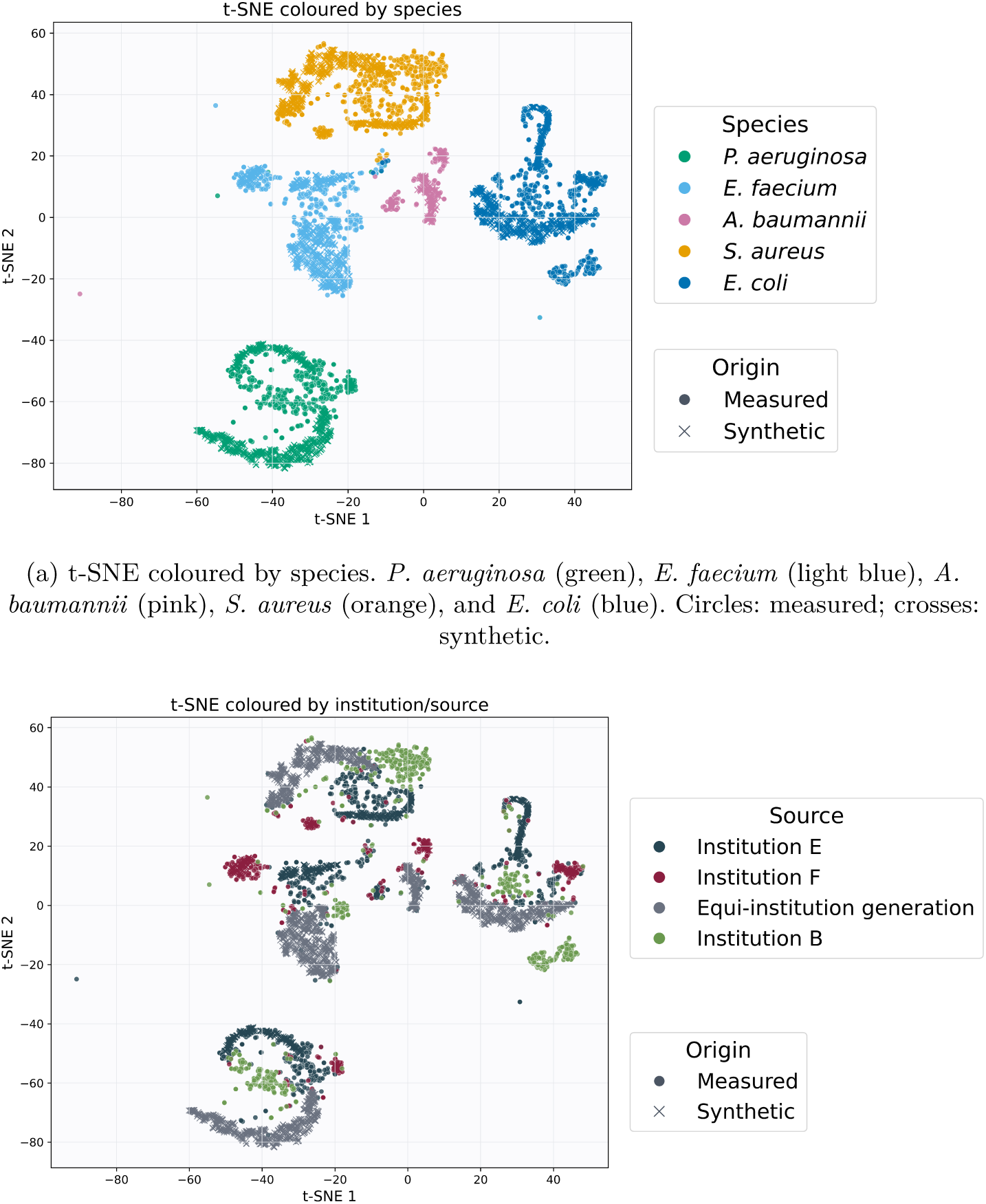
Latent-space organisation of measured and synthetic spectra. Real spectra are from the test set. **a,** Synthetic spectra retain species-associated neighbourhoods. **b,** Institution-conditioned generations follow institution-associated structure, whereas equi-institution sampling provides broader coverage of the within-species spectral space.

Together, these qualitative analyses suggest that the synthetic atlas preserves the main spectral patterns associated with each species. They also show that the generated spectra reflect differences between institutions. We next used quantitative metrics to evaluate these observations.

#### Quantitative validation of the synthetic spectra

We next quantified the properties of different synthetic data generated by different generative models to select one to construct the atlas.

We compared: a CBGM-VAE, a Generative Adversarial Network (GAN), a Diffusion Model, and a Variational Autoencoder (VAE). All models used the same backbone and reconstruction objective, selected through the ablation experiments reported in the Encoder and latent-size ablation section. CBGM-VAE was conditioned on both species and institution. The rest of the models were only conditioned on species. We also introduced a variant to the CBGM-VAE model, we called it **CBGM-VAE-O**. An orthogonality constrain was introduced between the representations of the species and the institution concept. This variant was designed to encourage a greater separation between taxonomic and acquisition-related information. For both CBGM models, we evaluated two generation strategies: (i) *equi-institution* sampling, in which we generate a sample to come equally probable from all institutions, and (ii) *institution-conditioned* sampling, in which a specific institution was explicitly imposed.

We evaluated the quality of the synthetic spectra based on distributional similarity, novelty, coverage of measured spectra, and preservation of species-discriminative information. Frechet MALDI distance (FMD) and Maximum Mean Discrepancy (MMD) measured distributional similarity to real spectra; novelty distance (ND) quantified similarity to same-species training examples; and coverage distance (CD) measured how closely synthetic spectra covered the measured samples. Finally, we trained a k-nearest-neighbour classifier on measured spectra and evaluated its macro-F1 on synthetic spectra as a measure of whether species-discriminative information was retained.

The Diffusion Model and GAN models reported the lowest species-discrimination power (Fig. 4a) and were rejected. In contrast, the CBGM-VAE models showed the best species classification (Fig. 4a) and the best trade-off between distributional similarity and species separability (Fig. 4b). Generating spectra for a specific institution gave lower FMD and MMD, meaning that the synthetic spectra more closely matched the measured training data. However, the kNN classified them less accurately. With equi-institution generation, the spectra were less closely matched to the training distribution but were more accurately classified by species. Therefore, we decided to use CBGM-VAE as the generator for the atlas.

**Fig. 4:**
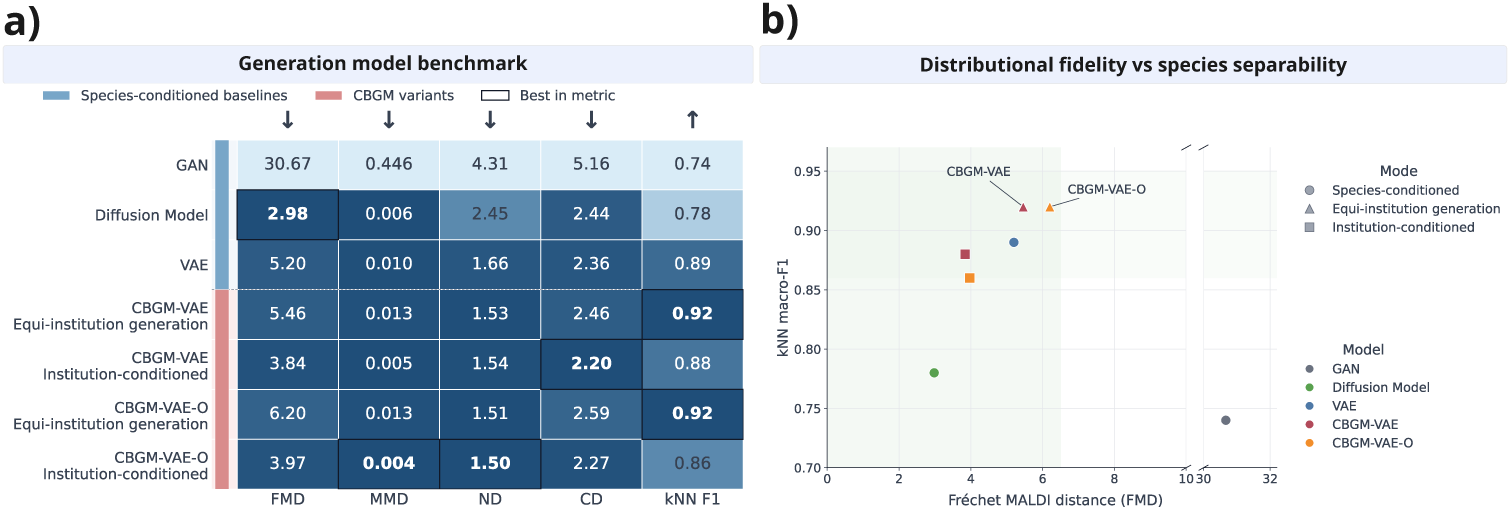
Quantitative validation and selection of the atlas generation strategy. **a,** Comparison of different generative models across distributional, coverage, and species-discriminative metrics. **b,** Trade-off between distributional fidelity and species separability across generative models and sampling strategies. Complete numerical results can be found in Supplementary Table S3. All metrics are defined in Evaluation and statistical analysis, and their mathematical formulation can be found in Evaluation metrics.

Generation quality varied across species. Some species with few measured training spectra showed lower kNN classification performance on synthetic data. This suggests that limited training data can affect generation quality. Further details are provided in the Supplementary Discrimination decays with data availability section.

Under equi-institution generation, both CBGM-VAE and CBGM-VAE-O achieved the highest kNN macro-F1 of 0.92. As a design choice, we selected CBGM-VAE-O to build the atlas. We hypothesised that the orthogonality constrain could produce a broader range of spectra within each species. Next, we tested whether these synthetic spectra could train classifiers that identify species in external real samples.

### The synthetic atlas is a useful training resource

We then expanded on the following important question: **can synthetic data be used to train bacterial classifiers?**

Using bacterial species classification as a benchmark, we compared three different ways of training such classifiers: using only real spectra (blue line), using only synthetic spectra (red line), and using both real and synthetic (green line), as depicted in Fig. 5. Details on the training methodology can be found in the Evaluation of the atlas as a training resource section.

**Fig. 5:**
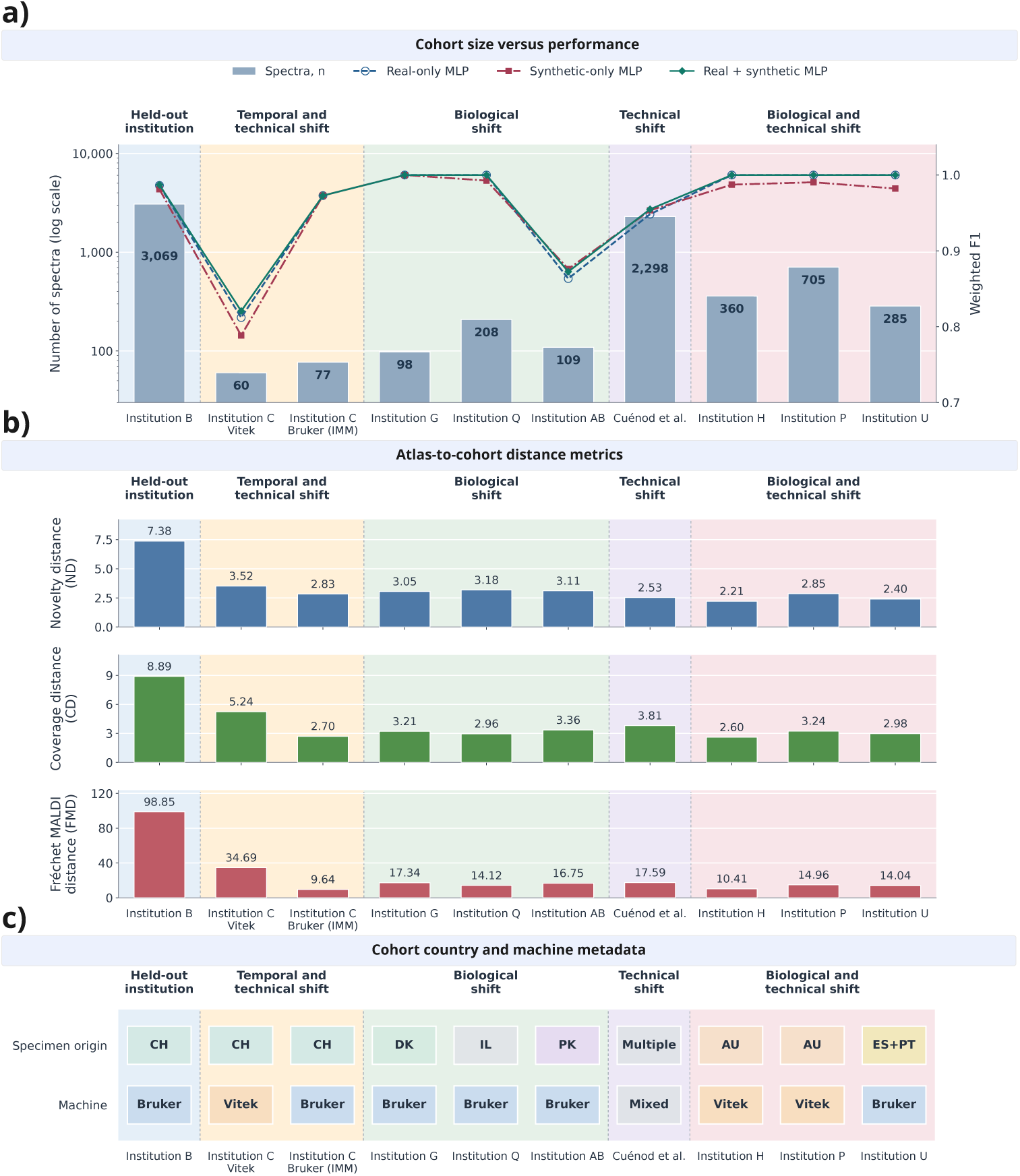
Synthetic atlas evaluation across real cohorts. **a,** Cohort size and species-identification performance of each training strategy. **b,** Atlas-to-cohort distance metrics across all cohorts. **c,** Country and machine metadata for each evaluated cohort. Numerical results are provided in Table S4.

We evaluated the three strategies on 7,269 real spectra from external cohorts. These cohorts covered five evaluation settings: (i) a held-out institution containing all 43 species, (ii) combined temporal and technical shift, (iii) biological shift, (iv) technical shift, and (v) combined biological and technical shift. Details on the methodology to group the shifts can be found in the External out-of-distribution cohorts section.

First, we found that classifiers trained exclusively on synthetic spectra achieved performance close to those trained on real data. On Institution B, synthetic-only training achieved a weighted F1 of 0.981, compared with 0.986 for real-only training. Similar results were observed across other evaluation settings. Under biological shift, both strategies achieved a pooled weighted F1 of 0.965. In the technical-shift cohort from Cúenod et al. [7], synthetic-only training even outperformed real-only training, reaching 0.953 compared with 0.948. These results show that the atlas preserves species information that can be learnt from synthetic spectra alone and transferred to real samples, including those from unseen institutions and acquisition settings.

Beyond its stand-alone use, the atlas also improved species classification when combined with real training data. This strategy achieved the highest weighted F1 across all five evaluation settings. On Institution B, weighted F1 increased to 0.987 when adding synthetic spectra. Similar modest improvements were observed under combined temporal and technical shift (0.903 to 0.906), biological shift (0.965 to 0.968), and technical shift (0.948 to 0.955).

We also measured how different the external cohorts were from the synthetic atlas using ND, CD, and FMD (Fig. 5b). Institution B showed the largest distances, followed by Institution C using VITEK. However, these distances did not follow a clear pattern across the different shift types.

Together, these results demonstrate the value of the synthetic atlas as a training resource. Classifiers trained only on synthetic spectra achieved performance close to those trained on real data, while combining both generally provided the best results. The atlas can therefore be used as a stand-alone training dataset or to complement existing measured data, providing a resource for developing MALDI–TOF MS machine-learning models.

To make the synthetic atlas easier to use, we developed an interactive web resource^1^ (Fig. 6) along with the release of the dataset [14]. Users can browse a lightweight preview of the atlas or upload their own MALDI–TOF MS for species identification. It is intended for research and benchmarking purposes. The database is not IVDR conform and should not be used for diagnostic decisions.

**Fig. 6:**
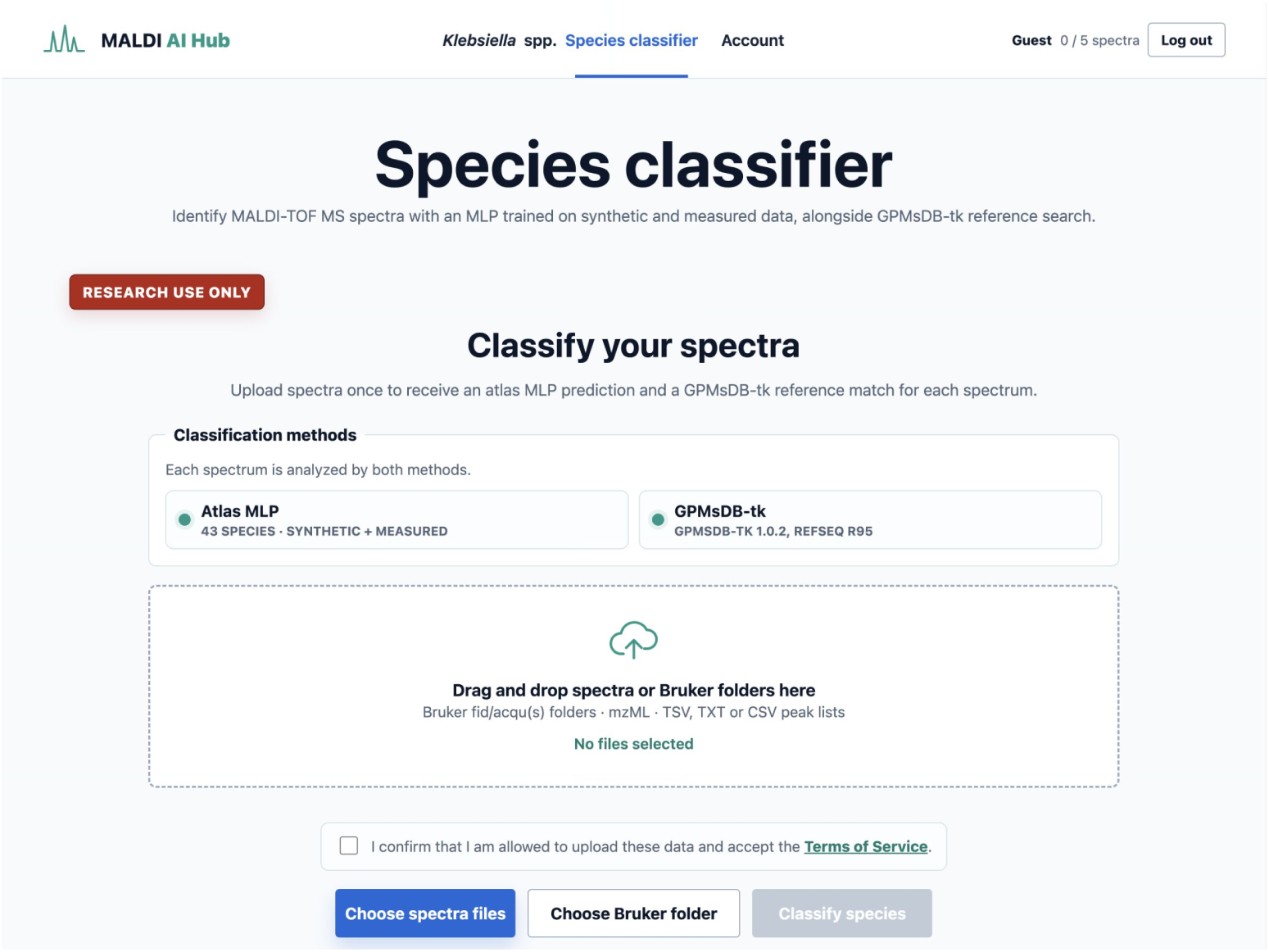
Interactive synthetic atlas resource available for research use only.

## Discussion

This work studies MALDI–TOF MS generation at a scale and evaluation setting closer to real deployment conditions. Building on concept-bottleneck generative modelling [17], we trained a generator on publicly available spectra from six institutions across three countries and 43 clinically relevant species [13]. Our results show that synthetic spectra preserve species-discriminative information and can be used to train classifiers that identify bacteria in real external data, achieving performance close to models trained on real spectra. Moreover, combining synthetic and real data improved species classification in several evaluation settings, while matching real-only performance in the others. These findings demonstrate that synthetic MALDI–TOF MS data can serve both as a stand-alone training resource and as a complement to measured data. We therefore release an open synthetic atlas containing 4.3 million spectra, with 100,000 spectra per species, for the research community [14].

Our results also highlight the importance of modelling acquisition context during synthetic generation. Previous MALDI–TOF MS augmentation approaches have shown benefits for resistance prediction [6, 10, 18] and ribotype classification [11], but do not explicitly account for the institution where the spectra were acquired. Without this information, models may be biased towards institutions contributing more spectra, potentially favouring their acquisition conditions and dominant bacterial lineages. We therefore modelled the institution explicitly and sampled uniformly across institutions to reduce the influence of any single source. We hypothesise that this approach can help capture species information shared between institutions while preserving broader within-species variation. Consistent with this hypothesis, equi-institution generation appeared less restricted to individual institution-associated regions in the t-SNE visualisation. We believe that it is a promising strategy for building more generalisable synthetic training resources.

Despite these results, several limitations remain. Although the atlas contains the same number of synthetic spectra per species, generating more samples cannot compensate for limited biological diversity in the original data. Additional real spectra are therefore still needed, particularly for under-represented species. Another limitation is that institution labels are only a proxy for acquisition conditions. Differences between institutions may reflect technical factors, such as instrument type, calibration, sample preparation, agar medium, culture conditions, or time between culture and spectral acquisition, but also biological differences between bacterial strains. These factors also contribute to interoperability challenges between MALDI–TOF MS platforms, where differences in acquisition, preprocessing, and data formats make it difficult to combine spectra from different manufacturers. Although the atlas supported species classification across multiple platforms, performance varied between cohorts and instruments. Future public datasets should therefore provide more diverse spectra, standardised data formats, detailed acquisition metadata, and quality-control information to support more reliable and generalisable synthetic resources.

In conclusion, this work provides a large-scale synthetic training resource for MALDI– TOF MS machine learning. Future work should extend the CBGM-VAE towards richer and conditioning schemes incorporating antimicrobial resistance, virulence, and bacterial typing labels. The released atlas should also be explored as a large-scale corpus for self-supervised pretraining of MALDI–TOF MS encoders [6, 10]. These applications will require further validation to determine whether synthetic spectra preserve the biological information needed beyond species identification. More broadly, our results suggest that synthetic and real MALDI–TOF MS data can complement each other in the development of machine-learning models. The atlas released here is therefore not an endpoint, but an open starting point for developing larger and more generalisable models for clinical microbiology.

## Methods

### Study design and data resources

#### Main training resources

We curated three public MALDI–TOF MS resources spanning six institutions across three countries: DRIAMS (Institutions A–D, Switzerland) [5], MALDI-TOF MS and Antimicrobial Resistance of Infectious Species at Marañón (MARISMA) (Institution E, Spain) [15], and University Medical Center Gö ttingen (UMG) (Institution F, Germany) [16]. We selected public databases containing at least 50,000 spectra to ensure sufficient data for training and to facilitate balanced contributions from each resource. All spectra were acquired using Bruker Daltonics instruments.

We harmonised species labels and institution metadata across the three resources and removed spectra that did not meet our quality-control criteria. The exclusion criteria and filtering procedures are described in Dataset curation and quality control. Species identification followed the procedures reported for each original dataset. In DRIAMS, identification was performed using the Microflex Biotyper Database (MBT 7854 MSP Library, BDAL V8.0.0.0 7311-7854, research-use only) with flexControl software (Bruker Daltonics, v3.4) [5]. In MARISMA, species were identified using the routine diagnostic database available each year, with an identification score above 1.7 [15]. In UMG, identification was performed using the MALDI Biotyper software suite (v4.1, database v16) [16].

#### External out-of-distribution cohorts

We assembled the external out-of-distribution panel from two sources: cohorts provided by collaborating institutions and publicly available datasets from previous studies. All institutions are renamed from Institution A to Institution AB. Their full names and corresponding identifiers are provided in Institution naming and standardisation.

First, collaborating institutions provided MALDI–TOF MS spectra from bacterial isolates collected across different geographical and clinical settings. Some cohorts also included isolates originating from additional institutions. For example, Institution U collected isolates from 17 institutions across Spain and Portugal, all measured using a single acquisition pipeline.

Second, we included two publicly available external datasets: the Institution H *Pseudomonas aeruginosa* cohort [8] and the multicentre cohort from Cúenod *et al.* [7]. The latter includes spectra collected across multiple laboratories before and after an intervention to improve and standardise MALDI–TOF MS acquisition procedures.

We grouped the external cohorts according to the main sources of distribution shift, as shown in Table 1:

**Table 1:** External out-of-distribution cohorts grouped by distribution shift. *n* denotes the number of spectra included in evaluation. “Orig. countries” and “Orig. institutions” refer to isolate origin, whereas “Acq. sites” and “Instr.” describe the acquisition sites and instruments used. Institution identifiers are defined in Institution naming and standardisation. The total row reports unique counts across all cohorts.

| Cohort | $n$ | Orig. countries | Orig. institutions | Acq. sites | Instr. | Scenario |
| --- | --- | --- | --- | --- | --- | --- |
| **Held-out institution** |  |  |  |  |  |  |
| Institution B | 3,069 | 1 | 1 | 1 | 1 | Unseen institution |
| **Temporal and technical shift** |  |  |  |  |  |  |
| Institution C | 137 | 1 | 1 | 2 | 2 | Temporal and dual-instrument acquisition |
| **Biological shift** |  |  |  |  |  |  |
| Institution G | 98 | 1 | 1 | 1 | 1 | Different isolate population, measured at the IMM |
| Institution Q | 208 | 1 | 1 | 1 | 1 | Different isolate population, measured at the IMM |
| Institution AB | 109 | 1 | 1 | 1 | 1 | Different isolate population, measured at the IMM |
| **Technical shift** |  |  |  |  |  |  |
| Cuénod *et al.* [7] | 2,298 | 12 | 35 | 35 | 3 | Multicentre acquisition variability |
| **Biological and technical shift** |  |  |  |  |  |  |
| Institution H [8] | 360 | 1 | 1 | 1 | 1 | Different isolate population and acquisition |
| Institution P | 705 | 1 | 1 | 1 | 1 | Different isolate population and acquisition |
| Institution U | 285 | 2 | 17 | 1 | 1 | Multi-origin, single-acquisition |

##### Held-out institution

Institution B was excluded from training and validation. This allowed us to evaluate the atlas in an unseen institution that contains all 43 bacterial species.

##### Temporal and technical shift

Institution C was represented in the training data, but the external samples were collected in 2025–2026, compared with the training spectra acquired in 2018. Moreover, the training spectra were acquired using an older Bruker instrument, while the external samples were measured using two different instruments: a VITEK system at Institution C and a Bruker system at the Institute for Medical Microbiology (IMM) at the University of Zürich, Switzerland. We therefore considered both temporal and technical differences. Although the isolates originated from an institution already represented in training, changes in bacterial populations over time cannot be excluded.

##### Biological shift

Institutions G, Q, and AB provided bacterial isolates from different countries, which were all measured at the IMM in Zürich using the same Bruker instrument. This allowed us to keep acquisition conditions consistent across these cohorts while evaluating differences in bacterial populations.

##### Technical shift

We used the multicentre dataset from Cúenod *et al.* [7]. In this study, the same bacterial isolates were measured across 36 laboratories in 12 countries. This allowed us to evaluate differences in instruments and acquisition procedures while keeping the bacterial samples consistent.

##### Biological and technical shift

Institutions H, P, and U provided isolates from different geographical origins. Unlike the biological shift cohorts, these spectra were acquired at their institution of origin. Therefore, these cohorts included differences in both bacterial populations and acquisition conditions.

These categories describe the main sources of variation in each cohort. They do not imply that biological and technical effects were completely independent.

#### Pathogen set

To focus on organisms of high clinical and public-health relevance, we based our species selection on the Lancet Global Burden of Disease Study [13]. Some bacterial groups in this study are defined at the genus level, such as *Citrobacter* spp. or other enterococci. Since MALDI–TOF MS datasets are typically annotated at the species level, we expanded these groups to include the corresponding species available in our public datasets. This resulted in a list of 43 bacterial species. The complete list and its correspondence to the original pathogen groups described in [13] are provided in Supplementary Table S5.

We then filtered all public MALDI–TOF MS datasets to retain only spectra belonging to these 43 species. As a result, the number of spectra included from each database may differ from the total number available in the original resource.

#### Split strategy

We divided the measured spectra into training, validation, and test sets. The evaluation included three settings: (i) temporal generalisation within the same institution, (ii) generalisation to a held-out institution, and (iii) generalisation to external cohorts with different biological and technical conditions. Validation data were used for model selection and early stopping, while test data were held out for final evaluation. When repeated measurements from the same isolate were available, all related spectra were assigned to the same split to avoid data leakage. The complete dataset composition is shown in Table 2.

**Table 2:**
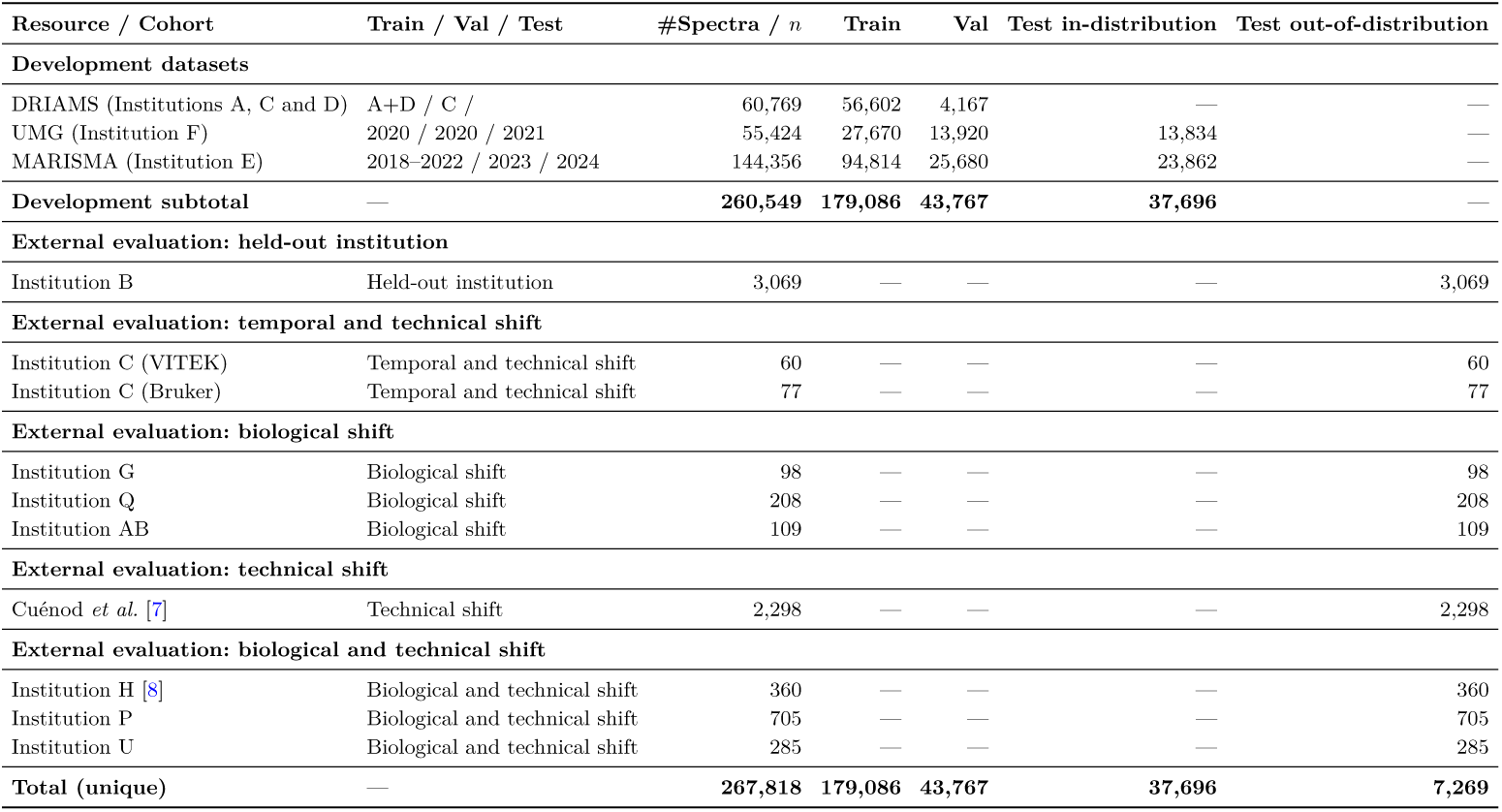
Dataset composition and evaluation splits. The upper section reports the development datasets after species filtering, including their training, validation, and test splits. The lower sections group the external evaluation cohorts by distribution shift, following Fig. 5. All external cohorts were excluded from model training and selection.

We used Institutions A and D for training, Institution C for validation, and Institution B as a held-out test institution. This allowed us to evaluate whether models trained on spectra from other institutions could identify species in an unseen institution.

For Institution E and Institution F, we used temporal splits to evaluate performance on future samples from the same institution. In Institution E, spectra from 2018– 2022 were used for training, 2023 for validation, and 2024 for testing. In Institution F, spectra acquired in 2020 were split into training (2/3) and validation (1/3) sets, while spectra from 2021 were reserved for testing. These splits allowed us to evaluate temporal changes while keeping the acquisition instrument unchanged.

Finally, we evaluated the atlas on external cohorts that were not used for training or model selection. These cohorts were grouped according to the main source of distribution shift: temporal and technical, biological, technical, or combined biological and technical shift. Together with the Institution B cohort, they formed the 7,269-spectrum external evaluation panel.

#### Spectrum representation and preprocessing

To combine spectra from different public databases, we represented all MALDI–TOF MS spectra on a common grid spanning *m/z* = 2,000–20,000, with 6,000 bins of width 3 Da. This representation matched the publicly available, pre-binned UMG data.

For all raw spectra, we applied the state-of-the-art preprocessing [5]: square-root transformation, Savitzky–Golay smoothing, SNIP baseline correction, and total-ion-current normalisation. We then restricted the spectra to the target mass range and summed intensities within each bin.

Finally, we applied the same transformations to all datasets: log_10_(1 + *x*) compression, removal of intensities below one standard deviation of the log-transformed spectrum, and per-spectrum min–max scaling to [0, 1]. Each spectrum was therefore represented as a normalised vector of 6,000 dimensions. Full preprocessing parameters are provided in the supplementary Preprocessing details.

#### Concept bottleneck generative model

To construct the synthetic atlas, we combined concept-bottleneck generative modelling (CBGM) [17] with a variational autoencoder (VAE) [19] and added a new orthogonality loss. The resulting model, CBGM-VAE-O, was designed to generate MALDI–TOF MS spectra conditioned on both bacterial species and acquisition institution. Fig. 7 illustrates the model architecture.

**Fig. 7:**
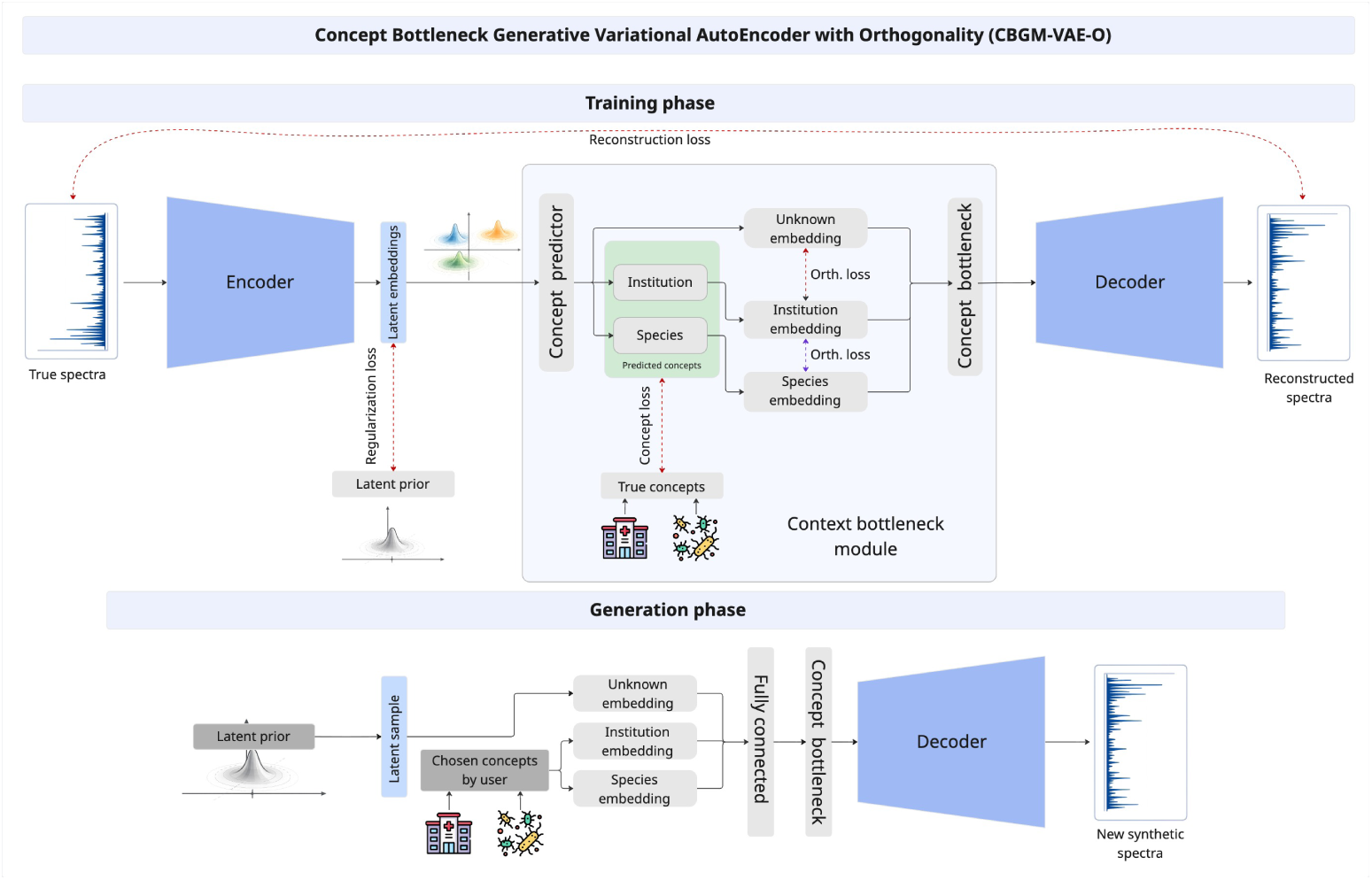
Architecture of the Concept Bottleneck Generative Variational Autoencoder with Orthogonality (CBGM-VAE-O). Top: During training, measured MALDI–TOF MS spectra are encoded into a latent representation. A concept predictor learns species and institution information, while an additional branch captures residual variation. The concept embeddings are combined through a bottleneck and passed to the decoder to reconstruct the input spectrum. The training objective includes reconstruction, latent regularisation, concept prediction, and orthogonality losses. **Bottom:** During generation, a latent sample is drawn from the learnt prior. Species and institution conditions are selected, combined with the residual embedding, and passed through the decoder to generate a new synthetic spectrum.

The encoder maps each input spectrum to a latent representation. This representation is then divided into three components: species, institution, and an unknown component that captures variation not explained by the first two. The components of species and institutions are learnt using their corresponding labels, while the unknown component is learnt without explicit supervision. All three components are combined through a concept bottleneck and passed to the decoder to reconstruct the input spectrum.

During training, species and institution labels supervise the corresponding concept predictors but are not directly provided to the decoder. The training objective combines reconstruction loss, variational regularisation, concept classification losses, and orthogonality constraints. These terms encourage the model to reconstruct spectra while learning the information associated with each concept.

We introduced an additional constraint between species and institution representations that is not canonical to the original formulation of the CBGM [17]. This constraint encourages the model to separate taxonomic and institution-associated information.

During generation, the model samples from the learnt latent distribution and allows the species and institution conditions to be selected. We evaluated two generation strategies: institution-conditioned generation, which specifies a single institution, and equi-institution generation, which samples institutions with equal probability.

The complete mathematical formulation, model architecture, and loss functions are provided in the supplementary Concept-Bottleneck VAE formulation.

#### Model selection

We trained all generative models using the Adam optimiser [20], with a learning rate of 10^−4^, a batch size of 128, gradient clipping at *ℓ*_2_ = 1.0, and early stopping based on the validation objective.

To reduce the effect of species imbalance, we used class-balanced sampling during training. At each epoch, we sampled up to 1,000 spectra per species, using replacement when fewer samples were available.

We compared two encoder–decoder architectures, a multilayer perceptron (MLP) and a one-dimensional convolutional neural network (1D-CNN), with latent dimensions of *z_i_* ∈ {64, 256, 768}. We also evaluated four reconstruction objectives: binary cross-entropy (BCE), focal loss, a balanced BCE–focal loss, and a hurdle-style loss [21] that separates peak presence from peak intensity.

Model selection was based on the balance between spectral fidelity, within-species variability, and preservation of species-discriminative information. The architecture and loss ablations are reported in the supplementary Loss ablation results. Full architecture definitions, training details, and loss functions are provided in the supplementary Training details, and Reconstruction losses sections, respectively.

#### Evaluation and statistical analysis

We evaluated the generated spectra using complementary metrics to measure distributional similarity, novelty, coverage of measured spectra, and preservation of species-discriminative information. Unless otherwise stated, all metrics were computed directly on the preprocessed MALDI–TOF MS spectra rather than on learnt embeddings.

To measure distributional similarity between real and synthetic spectra, we used Fŕechet MALDI distance (FMD), adapted from the Fŕechet Inception Distance [22], and Maximum Mean Discrepancy (MMD) [23]. Lower values indicate closer agreement between the two distributions.

We used novelty distance (ND) to measure the distance between each evaluated spectrum and its nearest real training spectrum of the same species. To assess how well synthetic spectra cover the measured data, we calculated coverage distance (CD). For each measured spectrum, we computed the Euclidean distance to its nearest synthetic spectrum and averaged these distances. Lower CD values indicate better coverage of the measured distribution by the synthetic data. For per-species evaluations, distances were calculated only between spectra belonging to the same species.

Finally, we evaluated whether the synthetic spectra retained species-discriminative information. We trained a *k*-nearest neighbours (kNN) classifier on real spectra and evaluated its macro-F1 on both measured test spectra and synthetic spectra. This allowed us to assess whether a classifier trained on measured data could recognise the species represented in the generated spectra.

Together, these metrics provided complementary measures of synthetic-spectrum quality. Their mathematical definitions and reporting conventions are provided in the Supplementary Evaluation metrics section.

#### Evaluation of the atlas as a training resource

To evaluate the atlas as a training resource, we compared three species-classification strategies: real-only, synthetic-only, and combined real and synthetic training. All strategies used the same multilayer perceptron (MLP) architecture, with 6,000 input features, four hidden layers of 1,024, 512, 256, and 128 units, ReLU activations, dropout of 0.1, and an output layer for the 43 bacterial species.

For real-only training, we sampled up to 1,000 spectra per species with replacement from the measured training set. This resulted in 43,000 training spectra. We reserved up to 128 spectra per species for validation from those, resulting in 5,504 validation spectra.

For synthetic-only training, we sampled 1,000 synthetic spectra per species for training without replacement. Then, we selected separate set of 128 spectra per species for validation. This resulted in 43,000 training spectra and 5,504 validation spectra, with equal representation of all 43 species.

#### For combined training, we combined both datasets

All classifiers were trained using Adam with a learning rate of 10^−3^, weight decay of 10^−4^, batch size 256, and gradient clipping at a norm of 1. We used cross-entropy without additional class-balancing during training. All experiments used random seed 42.

Models were trained for a maximum of 40 epochs, with early stopping after eight epochs without improvement in validation loss. The checkpoint with the lowest validation loss was selected for evaluation.

## Data availability

The public MALDI–TOF MS datasets used in this study are available from their original sources: DRIAMS [5], MARISMA [15], UMG [16], Alfred Hospital [8] and Cúenod *et al.* cohort [7]. The synthetic MALDI–TOF MS atlas generated in this work, comprising 4.3 million spectra, is available in [14]. The webapp tool is available^2^.

## Code availability

All code used in this study is available in GitLab^3^. The repository includes the implementation of the models evaluated in this work, as well as the scripts required for data preprocessing, model training, synthetic spectrum generation, and reproduction of the reported analyses and figures.

## Acknowledgements

A.G.-L. was supported by the European Union’s Horizon Europe research and innovation programme under the Marie Sk-lodowska-Curie Actions grant agreement No. 101278403. Views and opinions expressed are those of the author(s) only and do not necessarily reflect those of the European Union or the European Research Executive Agency. Neither the European Union nor the granting authority can be held responsible for them. A.E. was supported by the Swiss National Science Foundation (grant reference 213019), Helmut Horton Foundation, and an unrestricted grant by the University of Zurich. BPH and NLS are supported by Investigator Grants from the National Health and Medical Research Council, Australia (GNT1196103 to BPH, GNT2033803 to NLS). We thank Anna Diversi and Diana Albertos Torres for measuring the in-house MALDI-TOF MS at the Institute for Medical Microbiology, University of Zurich, and for their support with technical data collection.

## Author contributions

A.G.-L. developed the methodology, implemented the code, carried out the experiments, analyzed the data, and drafted the manuscript. E.M. and Y.G. contributed to interpreting the results, preparing figures, and discussing model design, loss selection, and the biological plausibility of the generated spectra. K.K., N.R., E.H., E.S.-C., B.P.H., K.K.S., M.A.S., M.O., J.M.-G., M.H.-G., and N.L.S. supported data collection and curation for their respective cohorts and reviewed the manuscript. A.E. supervised the project, provided resources and strategic guidance, and contributed to scientific discussions and interpretation of the results. All authors revised the manuscript and approved the final version.

## Competing interests

The authors declare no competing interests.

## Supplementary Results

### Public data clusters by institution

Fig. S1 shows the projection t-SNE of a random subset of the ESKAPEE pathogens found in the training set: *A. baumannii*, *E. asburiae*, *E. cloacae*, *E. kobei*, *E. faecium*, *E. coli*, *K. pneumoniae*, *P. aeruginosa*, and *S. aureus*.

These results support the hypothesis shown in An open synthetic MALDI–TOF MS atlas: the spectra are grouped mainly by country, database, species, and institution, which are confounded. However, from those categories, institution provides the most granular level of clustering as shown between Institutions A and D which are from the same country and database but still cluster by institution. Based on these results, we chose both species and institution as the main concepts to condition the generative model on.

### Qualitative comparison of real and synthetic spectra

This section complements the results shown in the main Qualitative comparison of real and synthetic spectra section. Fig. S2 shows a zoom in of Fig. 2 shown in the main document to help the reader focus on the details of the region between 6000 and 8000 m/z for five different pathogens: *A. baumannii*, *E. faecium*, *E. coli*, *P. aeruginosa*, and *S. aureus*.

**Fig. S1:**
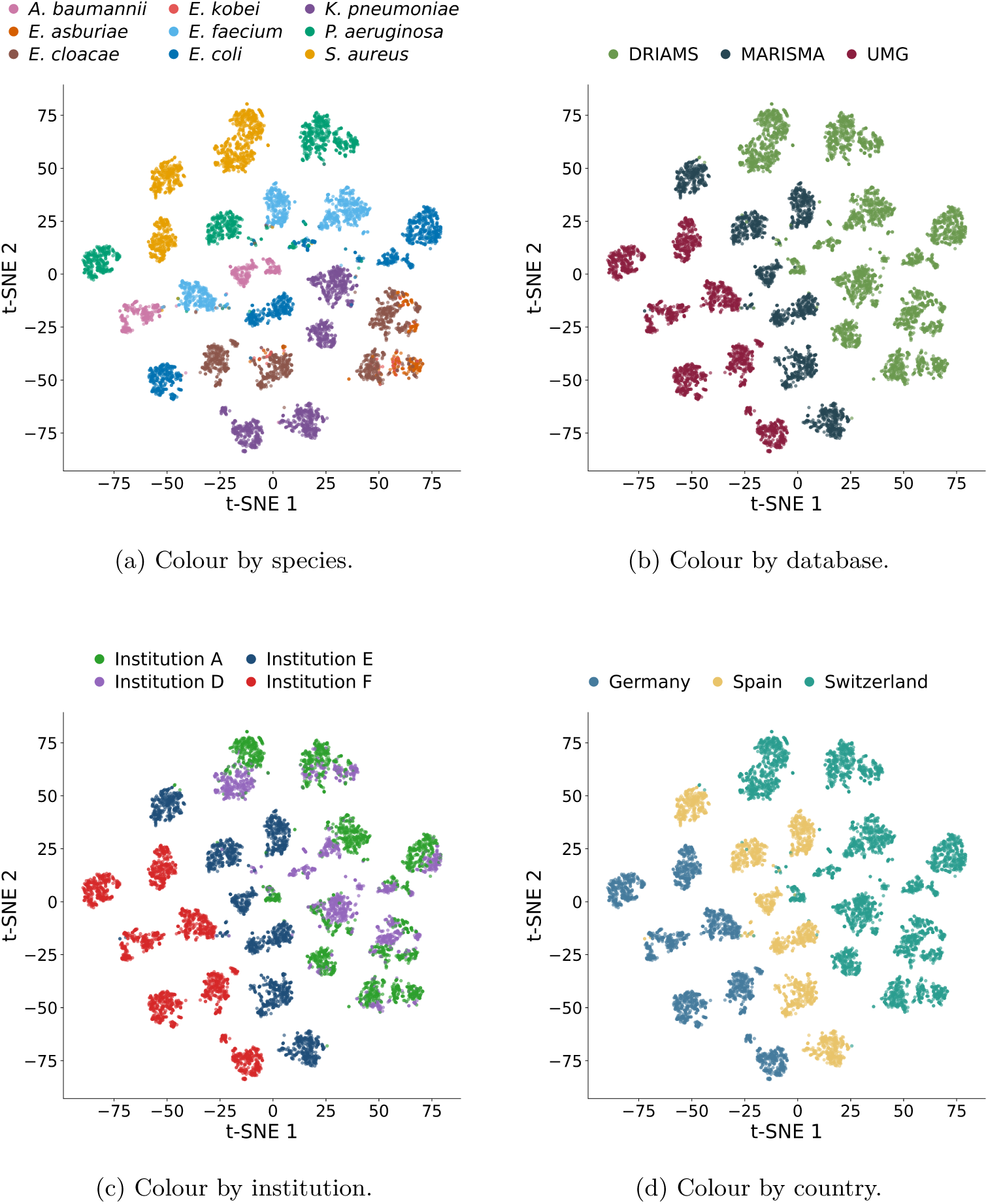
Metadata-associated structure in the harmonised MALDI–TOF MS data. The same t-SNE representation is coloured by (a) species, (b) database, (c) institution, and (d) country.

**Fig. S2:**
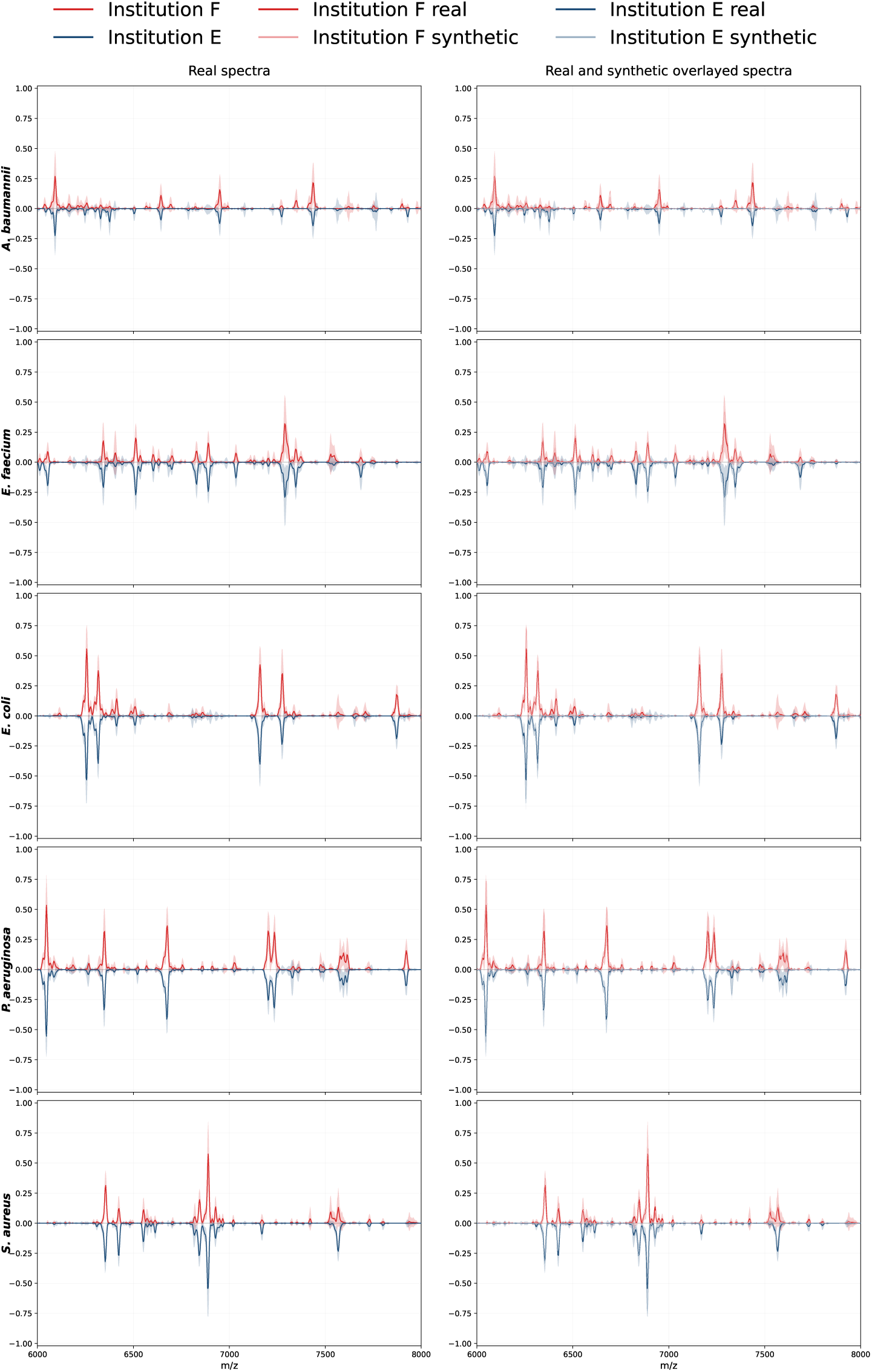
Zoom in to the region between 6000 and 8000 m/z.

### Discrimination decays with data availability

This analysis complements the results presented in Quantitative validation of the synthetic spectra. We investigated whether the number of measured training spectra per species affected the quality of the generated data. To assess this, we trained a kNN classifier on real spectra and evaluated its per-species balanced accuracy on both real Institution B spectra and synthetic spectra (Fig. S3).

**Fig. S3:**
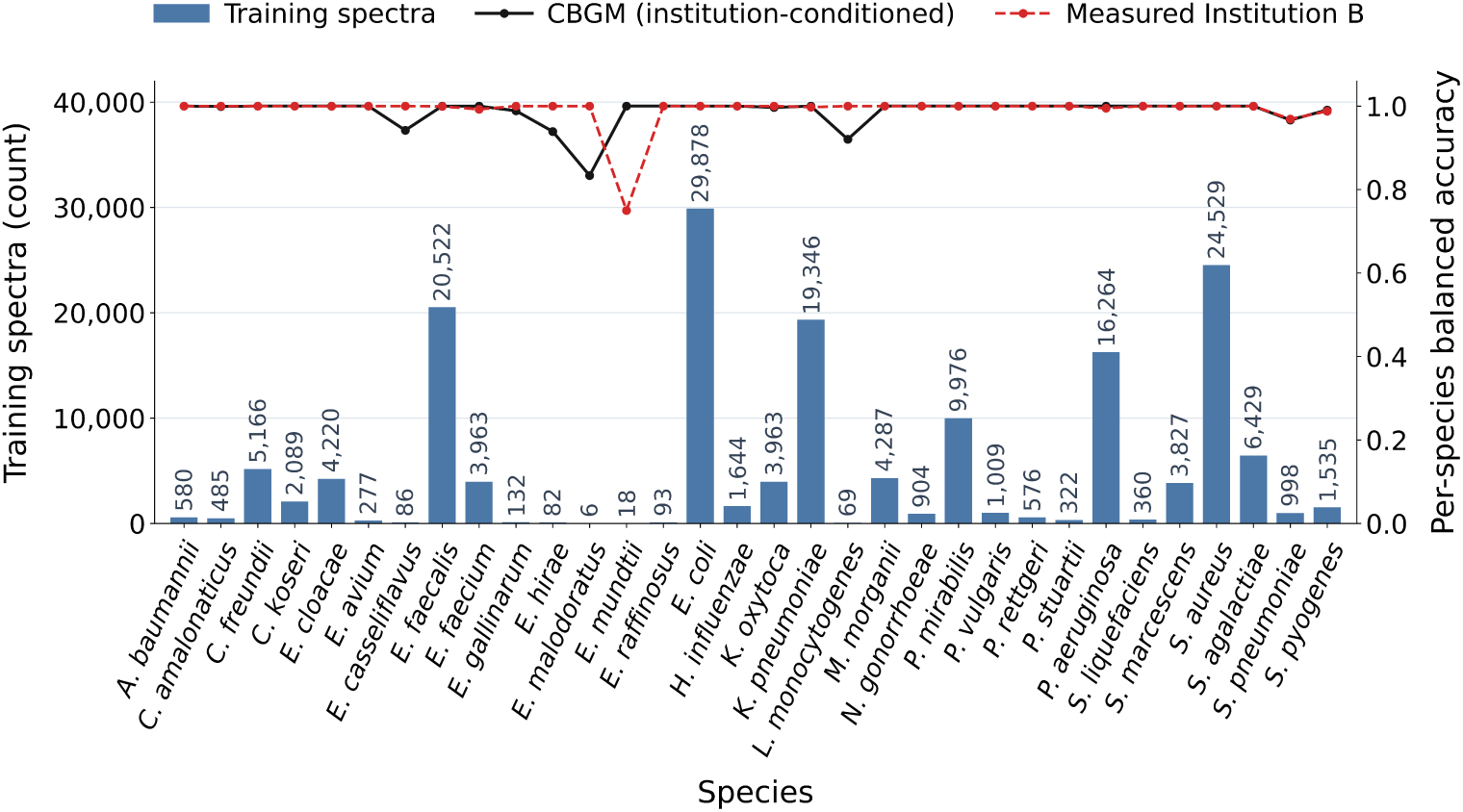
Effect of training data availability on synthetic-spectrum classification. Bars show the number of measured training spectra per species. The lines show per-species balanced accuracy of a kNN trained on real spectra and evaluated on real Institution B spectra (red) and synthetic spectra (black). The largest decreases in synthetic-spectrum classification occur among species with fewer than 150 measured training spectra.

The classifier achieved high balanced accuracy on real Institution B spectra and most synthetic species. However, some synthetic species were more difficult to classify, particularly *E. casseliflavus* (*n* = 86 in training), *E. gallinarum* (*n* = 132), *E. hirae* (*n* = 82), *E. malodoratus* (*n* = 6), and *L. monocytogenes* (*n* = 69). All these species had fewer than 150 measured training spectra. Importantly, not all species below this threshold showed reduced performance.

These results suggest that limited measured training data can make it more difficult for the generator to learn species-discriminative spectral patterns.

### Encoder and latent-size ablation

This section complements the Quantitative validation of the synthetic spectra section of the main manuscript.

We evaluated how the encoder–decoder architecture and latent dimensionality affected synthetic-spectrum quality. We compared CNN and MLP architectures with three latent dimensions (*z* = 64, 256, 768), using metrics of distributional similarity (FMD and MMD), novelty (ND), coverage (CD), and species classification (kNN macro-F1). Table S1 summarises the results.

**Table S1:**
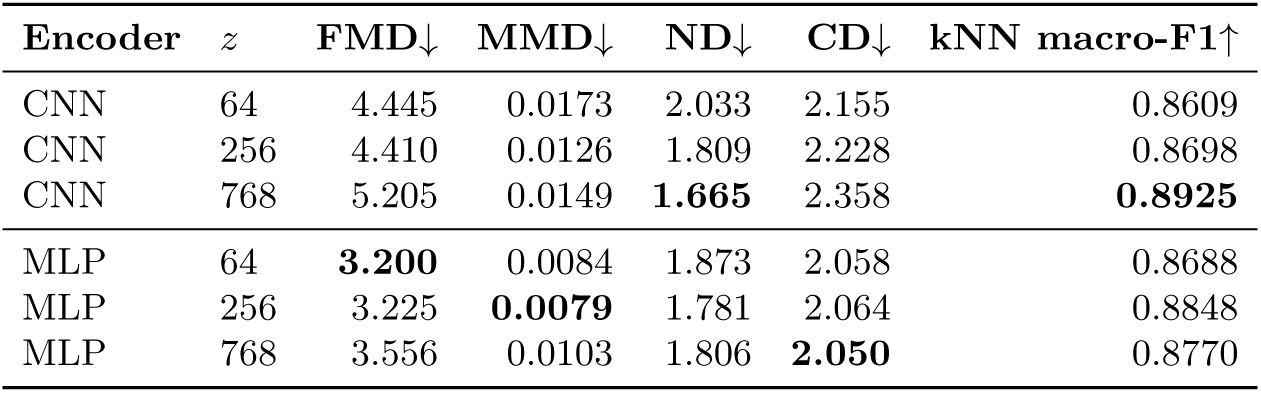
Encoder and latent-size ablation. Best values per metric are highlighted in **bold**. Lower values are preferred for FMD, MMD, ND, and CD, while higher values are preferred for kNN macro-F1.

No single configuration performed best across all metrics. The MLP configurations achieved the lowest FMD (*z* = 64), MMD (*z* = 256), and CD (*z* = 768), with each configuration leading in one metric. In contrast, the CNN with *z* = 768 achieved the lowest ND (1.665) and the highest kNN macro-F1 (0.8925), performing best in two of the five metrics.

We therefore selected **CNN with** *z* = 768 for the subsequent experiments. Although it did not provide the best distributional similarity or coverage, it achieved the strongest species-classification performance while keeping generated spectra close to the measured data. We prioritised these properties because the atlas was intended to preserve species information that could be used to train bacterial classifiers.

### Loss ablation

This section complements the Model selection section of the main manuscript.

With the backbone fixed to the selected CNN architecture (*z* = 768), we evaluated the effect of the reconstruction loss. Table S2 compares binary cross-entropy (BCE), focal loss, a balanced BCE–focal objective, and a hurdle-style loss designed to separate peak presence from peak intensity.

**Table S2:** Loss ablation for CNN, *z* = 768. Best value per metric is highlighted in bold. Values are rounded to two decimals.

| Encoder | $z$ | Loss | FMD↓ | MMD↓ | ND↓ | CD↓ | kNN F1↑ |
| --- | --- | --- | --- | --- | --- | --- | --- |
| CNN | 768 | Binary cross entropy | **4.03** | **0.01** | **1.72** | **2.18** | 0.87 |
| CNN | 768 | Focal | 7.82 | 0.05 | 2.57 | 2.76 | 0.88 |
| CNN | 768 | Balanced BCE–Focal ($\mathcal{L}_{BF}$) | 4.22 | **0.01** | **1.72** | 2.25 | **0.89** |
| CNN | 768 | Hurdle surrogate | 29.85 | 0.33 | 5.01 | 5.10 | 0.87 |

BCE and balanced BCE–focal achieved similar results and performed better than the other losses across most metrics (Table S2). BCE obtained the lowest FMD and CD, while both losses achieved comparable MMD and ND. Balanced BCE–focal achieved a slightly higher kNN macro-F1 (0.89 versus 0.87). However, this difference was small, and neither loss was consistently better across all metrics.

We selected **balanced BCE–focal** for the subsequent experiments. Our hypothesis was that combining both losses could preserve the good distributional similarity achieved by BCE while using the focal component to give more weight to difficult-to-reconstruct spectra. This could be particularly useful for species with more variable spectral patterns or limited training data.

### Numerical results of benchmarking

This section provides the numerical results supporting the generator selection described in Quantitative validation of the synthetic spectra section. Table S3 compares the generative models using metrics of distributional similarity (FMD and MMD), novelty (ND), coverage (CD), and species classification (kNN macro-F1).

**Table S3:** Comparison of generative models and sampling strategies. All models use the same selected backbone and reconstruction objective (CNN, *z* = 768, balanced BCE–focal loss). Lower values are preferred for FMD, MMD, ND, and CD, while higher values are preferred for kNN macro-F1.

| Model | Generation mode | FMD↓ | MMD↓ | ND↓ | CD↓ | kNN F1↑ |
| --- | --- | --- | --- | --- | --- | --- |
| GAN | Species-conditioned | 30.67 | 0.446 | 4.311 | 5.16 | 0.74 |
| Diffusion model | Species-conditioned | **2.98** | 0.006 | 2.45 | 2.44 | 0.78 |
| VAE | Species-conditioned | 5.20 | 0.010 | 1.66 | 2.36 | 0.89 |
| CBGM-VAE | Equi-institution | 5.46 | 0.013 | 1.53 | 2.46 | **0.92** |
| CBGM-VAE | Institution-conditioned | 3.84 | 0.005 | 1.54 | **2.20** | 0.88 |
| CBGM-VAE-O | Equi-institution | 6.20 | 0.013 | 1.51 | 2.59 | **0.92** |
| CBGM-VAE-O | Institution-conditioned | 3.97 | **0.004** | **1.50** | 2.27 | 0.86 |

### Extended synthetic-atlas evaluation on external cohorts

This section provides the numerical results supporting Fig. 5 of the main manuscript. Table S4 provides the corresponding cohort-level results and atlas-to-cohort distances.

**Table S4:** Species-classification performance and atlas-to-cohort distances across external cohorts.

| Cohort | Country | Machine | *n* | Species | Real only F1 | Synthetic atlas F1 | Real + synthetic F1 | ND | CD | FMD |
| --- | --- | --- | --- | --- | --- | --- | --- | --- | --- | --- |
| **Held-out institution** |  |  |  |  |  |  |  |  |  |  |
| Institution B | CH | Bruker | 3,069 | 43 | 0.986 | 0.981 | **0.987** | 7.38 | 8.89 | 98.85 |
| **Temporal and technical shift** |  |  |  |  |  |  |  |  |  |  |
| Institution C | CH | VITEK | 60 | 4 | 0.812 | 0.788 | **0.821** | 3.52 | 5.24 | 34.69 |
| Institution C | CH | Bruker | 77 | 4 | **0.973** | **0.973** | **0.973** | 2.83 | 2.70 | 9.64 |
| **Biological shift** |  |  |  |  |  |  |  |  |  |  |
| Institution G | DK | Bruker | 98 | 2 | **1.000** | **1.000** | **1.000** | 3.05 | 3.21 | 17.34 |
| Institution Q | IL | Bruker | 208 | 3 | **1.000** | 0.993 | **1.000** | 3.18 | 2.96 | 14.12 |
| Institution AB | PK | Bruker | 109 | 6 | 0.863 | **0.876** | 0.873 | 3.11 | 3.36 | 16.75 |
| **Technical shift** |  |  |  |  |  |  |  |  |  |  |
| Cuénod *et al.* [7] | Multiple | Mixed | 2,298 | 8 | 0.948 | 0.953 | **0.955** | 2.53 | 3.81 | 17.59 |
| **Biological and technical shift** |  |  |  |  |  |  |  |  |  |  |
| Institution H | AU | VITEK | 360 | 1 | **1.000** | 0.987 | **1.000** | 2.21 | 2.60 | 10.41 |
| Institution P | AU | VITEK | 705 | 4 | **1.000** | 0.990 | **1.000** | 2.85 | 3.24 | 14.96 |
| Institution U | ES+PT | Bruker | 285 | 1 | **1.000** | 0.982 | **1.000** | 2.40 | 2.98 | 14.04 |

## Supplementary Methods

### Dataset curation and quality control

Across all institutions, we first harmonised the metadata fields required for downstream analysis, including at minimum the species label and site identifier. Spectra with missing species or missing genus annotations were discarded. We additionally removed spectra with fewer than 100 non-zero intensity values, which we treated as failed or non-informative acquisitions.

The set of species used in this study is defined by expanding The Lancet Global Burden Disease [13] study list to the species level. As indicated in Table S5, some groups in [13] were defined at the genus level. Therefore, when expanded to all species found in the public databases of that genus, the list expanded to 43 bacterial species in total.

**Table S5:** Species used in this study, defined as species-level taxa derived from the bacterial-pathogen list proposed in The Lancet Global Burden Disease [13] study. The right column presents the 43 species found jointly in the training databases that correspond to the pathogen definition in the left column.

| Definition in [13] | Species included in this work |
| --- | --- |
| *Acinetobacter baumannii* | *A. baumannii* |
| *Aeromonas* spp. | *A. caviae* |
| *Campylobacter* spp. | *C. coli*; *C. jejuni*; *C. rectus*; *C. ureolyticus* |
| *Citrobacter* spp. | *C. amalonaticus*; *C. braakii*; *C. freundii*; *C. koseri* |
| *Enterobacter* spp. | *E. asburiae*; *E. cloacae*; *E. kobei* |
| *Enterococcus faecalis* | *E. faecalis* |
| *Enterococcus faecium* | *E. faecium* |
| Other enterococci | *E. avium*; *E. casseliflavus*; *E. gallinarum*; *E. hirae*; *E. malodoratus*; *E. mundtii*; *E. raffinosus* |
| *Escherichia coli* | *E. coli* |
| *Haemophilus influenzae* | *H. influenzae* |
| *Klebsiella pneumoniae* | *K. pneumoniae* |
| Other *Klebsiella* species | *K. oxytoca*; *K. variicola* |
| *Legionella* spp. | *L. pneumophila* |
| *Listeria monocytogenes* | *L. monocytogenes* |
| *Morganella* spp. | *M. morganii* |
| *Neisseria gonorrhoeae* | *N. gonorrhoeae* |
| *Proteus* spp. | *P. hauseri*; *P. mirabilis*; *P. vulgaris* |
| *Providencia* spp. | *P. rettgeri*; *P. stuartii* |
| *Pseudomonas aeruginosa* | *P. aeruginosa* |
| *Serratia* spp. | *S. liquefaciens*; *S. marcescens* |
| *Staphylococcus aureus* | *S. aureus* |
| Group A *Streptococcus* | *S. pyogenes* |
| Group B *Streptococcus* | *S. agalactiae* |
| *Streptococcus pneumoniae* | *S. pneumoniae* |

### Institution naming and standardisation

To simplify the presentation of results and ensure consistent naming throughout the study, we assigned each institution a unique identifier, ranging from Institution A to Institution AB. These identifiers are used throughout the main text, figures, and supplementary materials instead of the full institution names. Table S6 provides the corresponding institution names and their assigned identifiers.

**Table S6:** Institution identifiers used throughout the study. Mapping between standardised institution labels (A–AB), their full names, countries, and ISO 3166-1 alpha-2 country codes.

| Institution | Name | Country | ISO code |
| --- | --- | --- | --- |
| Institution A | DRIAMS-A Hospital Basel | Switzerland | CH |
| Institution B | DRIAMS-B Canton Hospital Basel-Land | Switzerland | CH |
| Institution C | Kantonsspital Aarau | Switzerland | CH |
| Institution D | DRIAMS-D Laboratory service Viollier | Switzerland | CH |
| Institution E | Hospital General Universitario Gregorio Marañón | Spain | ES |
| Institution F | University Medical Center Göttingen | Germany | DE |
| Institution G | Aalborg Universitetshospital | Denmark | DK |
| Institution H | Alfred Hospital | Australia | AU |
| Institution I | Centro Hospitalar de Lisboa Ocidental | Portugal | PT |
| Institution J | Centro Hospitalar Lisboa Norte | Portugal | PT |
| Institution K | Centro Hospitalar Universitário do Porto | Portugal | PT |
| Institution L | Centro Hospitalar Universitário Lisboa Central | Portugal | PT |
| Institution M | Centro Hospitalar Universitário São João | Portugal | PT |
| Institution N | CHUA – Unidade de Portimão | Portugal | PT |
| Institution O | Consorcio Hospital General Universitario de Valencia | Spain | ES |
| Institution P | Doherty Institute | Australia | AU |
| Institution Q | Hadassah Medical Center | Israel | IL |
| Institution R | Hospital Clinic Barcelona | Spain | ES |
| Institution S | Hospital Garcia de Orta, Almada | Portugal | PT |
| Institution T | Hospital Infante Dom Pedro, Aveiro | Portugal | PT |
| Institution U | Hospital Ramón y Cajal | Spain | ES |
| Institution V | Hospital Reina Sofía | Spain | ES |
| Institution W | Hospital Universitario de A Coruña | Spain | ES |
| Institution X | Hospital Universitario Marqués de Valdecilla | Spain | ES |
| Institution Y | Hospital Universitario Son Espases | Spain | ES |
| Institution Z | Hospital Universitario Virgen Macarena | Spain | ES |
| Institution AA | Unidade Local de Saúde de Matosinhos | Portugal | PT |
| Institution AB | University of Haripur | Pakistan | PK |

### Preprocessing details

Raw spectra from DRIAMS and MARISMA spectra were processed in the following order:

1. Square-root transformation of non-negative intensities.
2. Savitzky–Golay [24] smoothing with a half-window of 10 (21 points) and polynomial order 3.
3. SNIP (Statistics-sensitive Non-linear Iterative Peak-clipping) [25] baseline correction with 20 iterations.
4. Total-ion-current normalization, dividing each intensity by the sum of intensities in the baseline-corrected spectrum.
5. Trimming to *m/z* = 2,000–20,000.
6. Binning into 6,000 bins of width 3 Da, using the sum of intensities within each bin.
7. log_10_(1 + *x*) compression.
8. Per-spectrum standard-deviation thresholding with threshold factor *α* = 1.
9. Per-spectrum min–max scaling to [0, 1].

UMG spectra were provided as pre-binned vectors on the same grid. We therefore applied only steps 7–9 to these vectors.

Step 7, in detail, means: for a binned intensity vector *x*, log compression gives

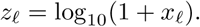

Then, in step 8, we compute the population standard deviation *σ*(*z*) across all 6,000 bins and apply

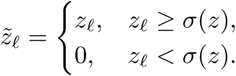

Finally, in step 9, we scale each spectrum as

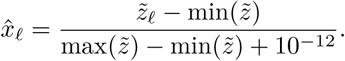

Vectors with no intensity variation are set to zero.

### Concept-Bottleneck VAE formulation

Given an input MALDI–TOF spectrum **x** ∈ R*^D^*, i.e., *D* = 6000, the encoder parameterizes a diagonal Gaussian posterior

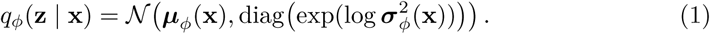

where 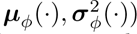 are the encoder network.

A latent sample is then obtained using the reparameterization trick:

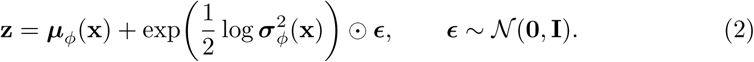

The sampled latent vector is not passed directly to the decoder. Instead, it is first mapped into a concept bottleneck representation that separates interpretable information from residual variation.

For each concept *k* ∈ {species, hospital}, the model applies a linear classifier to the latent sample **z**,

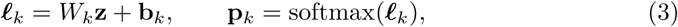

where ***ℓ****_k_* ∈ R*^Ck^* are the class logits, *C_k_* is the number of classes for concept *k*, and **p***_k_* is the corresponding categorical distribution. Rather than decoding a hard class label, the model uses a learnt embedding table

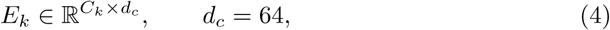

and computes the concept vector as the expected embedding under the predicted class distribution,

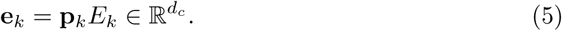

This construction is fully differentiable: if the target concept is known during conditional generation, one can simply replace **p***_k_*by a one-hot vector; otherwise, the model uses its own predicted probabilities.

In parallel, the model learns an “unknown” residual branch that captures variation not explained by species or hospital. This branch is obtained by a linear projection of **z** followed by dropout,

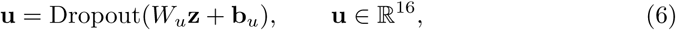

with dropout probability 0.2. The purpose of this branch is to retain nuisance or fine-grained information, such as strain-level variation, instrument effects, or other spectral factors that are not represented by the explicit concept variables. The dropout is added to ensure that the model does not rely completely on this branch and avoid using the known concepts.

The concept vectors and the unknown branch are then concatenated and passed through a bottleneck multilayer perceptron,

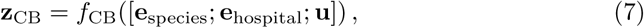

where [·; ·; ·] denotes concatenation. In the final model, this bottleneck maps a 64 + 64 + 16 = 144 dimensional input to a 768-dimensional representation. Thus, the decoder does not receive the raw latent vector **z**, but the concept-aware bottleneck vector **z**_CB_.

The decoder mirrors the encoder. It takes **z**_CB_ as input and maps it back to a length-6000 spectrum. In the CNN version used for the final model, the decoder first expands the bottleneck through fully connected layers and then reshapes the result into a low-resolution feature map. This feature map is progressively upsampled with one-dimensional convolutional blocks until the original spectral resolution is recovered. The decoder outputs a vector of logits **x**^ ∈ R^6000^. Since the reconstruction likelihood is Bernoulli, these logits are interpreted as unnormalized Bernoulli parameters; sigmoid is applied only when probabilities are needed for inspection or sampling.

Formally, the reconstruction term is

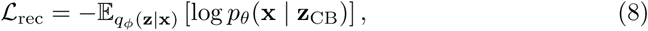

approximated with a single reparameterized sample of **z**. In the final model, the Bernoulli reconstruction loss was implemented as a weighted mixture of binary cross-entropy and focal loss on the decoder logits.

The prior over the latent space is a learnable diagonal Gaussian [26],

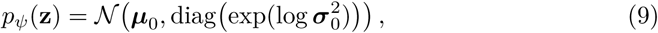

with trainable mean ***µ***_0_ and log-variance log ***σ***^2^ shared across all samples. This is more flexible than a fixed standard normal prior and allows the latent distribution to adapt to the training data. The KL term is computed between the encoder posterior and this learnable prior, with clipping of the encoder and prior log-variances to improve numerical stability.

The concept heads are trained with standard cross-entropy losses,

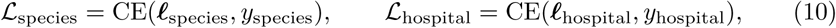

so that the latent representation predicts the annotated biological and acquisition-related concepts. In addition, an orthogonality regularizer encourages the unknown residual branch to be decorrelated from the concept embeddings,

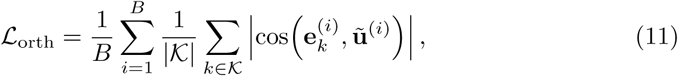

where **u**° denotes the unknown branch projected to the concept embedding dimension when necessary, and K is the set of known concepts. This penalty encourages the concept vectors and the unknown branch to represent different factors of variation.

An additional concept-concept orthogonality term can also be enabled to reduce redundancy between species and hospital embeddings.

Putting these components together, the total objective is

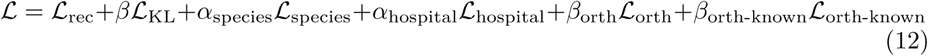

At generation time, the model can either infer the concept vectors from a sampled latent code or condition them explicitly. If a species or hospital is provided, the corresponding concept distribution is set to the one-hot label vector; otherwise, the model predicts the concept probabilities from **z**. In both cases, the unknown branch is still generated from the sampled latent code, and the concatenated bottleneck vector is decoded into a synthetic MALDI–TOF spectrum.

#### MLP variants

For the multilayer perceptron ablations, the encoder used layer widths proportional to latent size *z*,

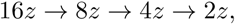

and the decoder used

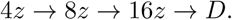

For example, with *z* = 768, the encoder widths are 12288 → 6144 → 3072 → 1536 and the decoder widths are 3072 → 6144 → 12288 → 6000.

### Training details

All models were optimized with Adam using a learning rate 1 × 10^−4^ and a batch size of 128 for up to 500 epochs. We applied gradient clipping with maximum *ℓ*_2_ norm 1.0 and used early stopping based on the validation objective with patience 50 epochs. Unless otherwise stated, experiments used random seed 42.

To mitigate species imbalance, we used class-balanced sampling at the epoch level. For each epoch, we drew up to 1,000 spectra per species for both training and validation. When fewer than 1,000 spectra were available for a species, sampling was performed with replacement; otherwise, sampling was performed without replacement.

We compared two encoder–decoder backbone families: (i) a multilayer perceptron and (ii) a one-dimensional convolutional neural network. For each family, we evaluated latent dimensionalities *z* ∈ {64, 256, 768}. The final model configuration was selected based on the validation ablation described in the Results.

Training was performed on a single H100 GPU. The final model was selected using the best validation performance.

### Reconstruction losses

#### Bernoulli likelihood and BCE

For normalized spectra *x* ∈ [0, 1]*^D^*, we first considered an element-wise Bernoulli likelihood parameterized by decoder logits. Under conditional independence across bins, maximizing the likelihood corresponds to minimizing the binary cross-entropy:

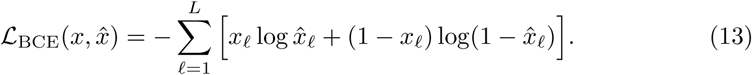

##### Balanced BCE–focal objective

For the final selected CBGM-VAE, reconstruction used a balanced BCE–focal loss:

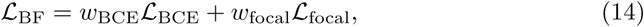

with *w*_BCE_ = 0.5 and *w*_focal_ = 0.5. The focal component used *γ*_+_ = 1.0, *γ*_−_ = 2.0, *α*_+_ = 1.0, and *α*_−_ = 1.0.

For each spectral bin *d*, let *x*^*_d_* denote the decoder logit and *p_d_* = *σ*(*x*^*_d_*) the corresponding Bernoulli probability, where *y_d_* ∈ {0, 1} is the binary target. The focal loss is defined elementwise by

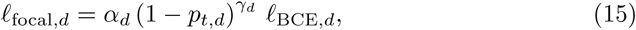

where

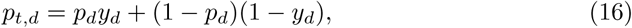

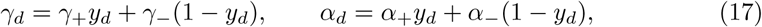

and

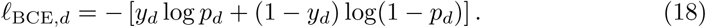

Equivalently, the focal term downweights easy examples through the factor (1 − *p_t,d_*)*^γd^* : when the model assigns high probability to the correct label, *p_t,d_* ≈ 1 and the loss contribution is strongly reduced, whereas misclassified or uncertain bins contribute more heavily. In our implementation, the reconstruction loss is obtained by summing the elementwise terms over all *D* bins, and the balanced objective averages the binary cross-entropy and focal components with equal weights.

##### Hurdle-style reconstruction objective

We also evaluated a hurdle-style objective that decouples peak presence from peak intensity. Using threshold *τ* = 0.05, we first define a binary peak mask

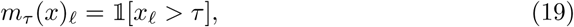

which defines where is a peak in the spectra. We then combine a presence term and an intensity term:

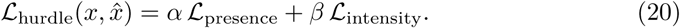

In code, these correspond to presence weight, which follows the BCE-focal objective and intensity weight which is a simple mean square error, respectively. The presence term weight was fixed to 1.0, while the intensity term weight was swept over {5, 10, 100}.

### Evaluation metrics

Unless otherwise stated, all evaluation metrics were computed directly in the preprocessed MALDI–TOF input space.

#### Fŕechet MALDI distance (FMD)

Let *µ_r_,* Σ*_r_* and *µ_g_,* Σ*_g_* denote the empirical means and covariance matrices of the real and generated spectra, respectively. We define

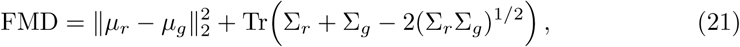

with lower values indicating closer distributional alignment.

#### Maximum mean discrepancy (MMD)

Using a radial basis function kernel *k*(·, ·), we compute

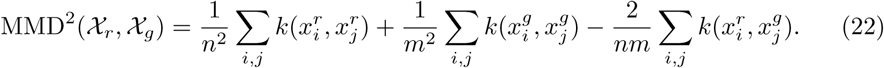

Lower values indicate closer alignment. We used the same kernel and bandwidth selection across all comparisons and interpret MMD comparatively rather than in absolute terms.

#### Novelty distance (ND)

For an evaluated sample **x** ∈ R*^D^* with class label *y*, let

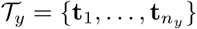

denote the set of training spectra belonging to the same class *y*. The novelty distance of **x** is defined as the Euclidean distance to its nearest same-class training neighbor:

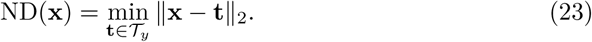

We report the mean ND over the evaluated set,

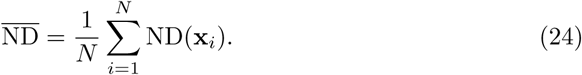

Lower values indicate that a sample lies closer to the observed training distribution of its class. For synthetic spectra, very low ND can indicate near-duplication or memorization2 of training examples, whereas high ND suggests that the generated spectrum is farther from the training distribution and may be more off-manifold.

##### Coverage distance (CD)

Coverage distance measures how closely the synthetic spectra cover the measured data. Let *R* = {*r*_1_*, …, r_N_*} and *S* = {*s*_1_*, …, s_M_*} denote the sets of measured and synthetic spectra, respectively. We define CD as

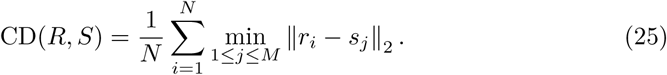

For each measured spectrum, we calculate the Euclidean distance to its nearest synthetic spectrum and average these distances. Lower CD values indicate that the synthetic spectra provide closer coverage of the measured samples.

For overall cohort evaluations, distances are calculated using pooled spectra. For per-species evaluations, nearest-neighbour searches are restricted to spectra of the same species.

#### kNN probe

To assess class-conditional consistency, we trained a *k*-nearest neighbors classifier with *k* = 5, uniform weights, Euclidean distance, and n jobs=-1 on real spectra and evaluated macro-F1 on both held-out real cohorts and generated samples. No hyperparameter tuning was performed for this probe.

#### Reporting conventions

Metric directions are indicated with arrows (↓ lower is better and **↑** higher is better).

## Footnotes

1 http://130.60.9.65/tools/synthetic-atlas

2 http://130.60.9.65/tools/synthetic-atlas

3 https://gitlab.uzh.ch/appliedmicrobiologyresearch/amr_publications/synth-maldi-atlas

## Notes

http://130.60.9.65/tools/synthetic-atlas

### Competing Interest Statement

The authors have declared no competing interest.

